# Switchable amplification of fluoresence from a photosynthetic microbe

**DOI:** 10.1101/167122

**Authors:** Anirban Bose, Sufi O Raja, Rajdeep Chowdhury, Somen Nandi, Sanhita Ray, Kankan Bhattacharyya, Anjan Kr Dasgupta

## Abstract

One known attribute of the photosynthetic apparatus is photon capture and generation of metabolic energy. The thermodynamic implications of fluorescence, invariably associated with the photosynthetic components is however poorly understood. In this paper we report a density dependent amplification of such fluorescence which can be interpreted as a thermodynamic strategy of controlled energy release by the cell. We show in support of this hypothesis that prolonged photo-exitation of cell free extract of *Rhodobacter capsulatus* SB1003 at 395 nm, induces fluorescence emission amplifying with time as long as the fluorophore density is above a critical level. The fact that the amplification disappears at low temperature and at dilute condition, is in accordance with the thermodynamic interpretation that energy is released as per requirement. Live cell imaging is also validation of the phenomenon even at the cellular level. Single cells of *Rhodobacter capsulatus* SB1003 shows time dependent loss of fluorescence, the process being reversed for cellular clusters. To explain the mechanism of this bistable fluorescence (F) amplification, variation of the scale free kinetic constant k=1/F (dF/dt) is studied at varying temperatures in presence and absence of static magnetic field. The sign of k shifts from positive to negative if T is lowered or if the system is diluted. But at low T, k again switches from negative to positive value, if static magnetic field is applied. The chain of events can be explained by a simple model assuming excretion of a porphyrin by the microbe and possible photon dependent aggregation behavior of such porphyrin complex, differential temperature and magnetic field sensitivity of the monomeric or aggregated forms of porphyrin being reported earlier.

## Introduction

Photosynthesis as Schrodinger’s description is the pivotal source of the negative entropy generation. While the conversion of photonic energy to proton motive force is a part of energy utilization the question remains as the regarding the functional significance of fluorescence always associated with the photosynthetic machinery. Thermodynamically fluorescence can be described as absorption of a higher photonic energy and release of photonic energy at lower wavelength. It is a non-thermal way of releasing the residual energy that is not converted to the useful metabolic work. This in turn may be useful in maintaining the thermostasis at the post photon-capture state.

The term bistability, is a concept implying a duality of steady states, pouring in biological literature in different contexts [1]. Broadly it expresses a switching from one steady state to another if a parameter is varied. Thus in genetics, epigenetics [2]and microbial ecology [3] bistability appears in order to describe existence of a dual genomic or ecological equilibria. It also appeared in sudden switching of oxygen concentration during the evolution of the bio-geosphere [4]. There was obviously a sudden surge of photosynthetic activity. Bistability in photochemical processes have been reported earlier in several photo-chromic compounds, polymers and derivatives [5, 6, 7, 8].

The term bistability is brought in this report in a context that is related to thermodynamic description. The cell may need the thermostatis when the photon capture centres have high density and vice versa. It therefore increases fluorescence emission with time. Similarly, at lower temparture one does not need any additional energy release mechanism.

We show bistability in fluorescence emission from *Rhodobacter capsulatus* SB1003, the switchability interestingly depends on fluorophore concentration and temperature. The bistability that we report here is not in terms of onset of fluorescence but in terms of amplification or quenching of the same.

Notably in spite of the presence of such a vast literature bistability has not been reported for a natural photosynthetic or cyno-bacteria. In general we find different classes of fluorescent biomolecules that are [9, 10] mostly susceptible to photo-bleaching. The work was inspired by our preliminary observation that we made while studying the fluorescence recovery after photobleaching (FRAP) experiment [11]. FRAP is based on the high propensity of bleaching by fluorophores and its recovery by diffusion of unbleached molecules. Our study shows the fluorescence from *Rhodobacter capsulatus* SB1003 became brighter instead of bleaching, under a confocal microscope during relatively prolong UV excitation.

Photosynthetic purple non-sulfur bacteria *Rhodobacter capsulatus* SB1003 is one of the most primitive photoheterotroph that contain photoreceptors in the form of porphyrin derivatives like non-covalently bound cofactors, bacterio-chlorophyll and bacteriopheophytin in the reaction centres of their photosynthetic membrane complexes [12]. This class of α-Proteobacteria has been employed as a model system for studying photosynthesis [13] and different classes of porphyrin production under various growth conditions [14]. Therefore bistability of the photosynthetic machinery of this purple bacteria seems to have important ecological and evolutionary implications.

These bacteria possess behavioral competencies to respond external stimuli by altered electron transport system which modulate their intra-cellular energy level to obtain optimal metabolic activity known as energy-taxis [15]. Such behavioral response is very essential survival strategy for obtaining highest metabolic yields at different external energy levels. The photosynthetic microbes are of particular interest as they induce a thermodynamically favorable condition for supporting the ecological network of different life forms by contributing negative entropy in the form of photon capture [16]. The general conjecture is that a sensory perception for species like *Rhodobacter* can respond to external light stimulus through electron transport system in their photosynthetic apparatus [17]. Numerous models have been reported where the role of microbes in overall metabolic energy transfer circuitry of different ecological niche have been emphasized [18, 19, 20, 21].

Notable point that we report in the present paper is that spin perturbations agents like static magnetic field [22, 23, 24] can play important role in this bistable emission. Possible roles of porphyrin aggregation, their excited state photo chemistry including radical pair formation [25, 26] and alteration of singlet-triplet ratio [27, 28] has been studied to understand the regulation process for amplified fluorescence emission. In addition temperature, concentration and interaction with static magnetic field effect have been investigated that could help one to truly embed the photon utilization and regulation machinery in designing and device level implementation of artificial photosynthesis.

## Material and Methods

All chemical reagents were used of analytical grade Sigma-Aldrich (USA) or SRL (India) products.

### Bacterial Growth

The bacteria *Rhodobacter capsulatus* strain SB1003 was a gift from Dr. Patrick Hallenbeck, Department of Microbiology and Immunology, University of Montreal, Canada. For experimental purposes *Rhodobacter capsulatus* SB1003 were grown photoheterotrophicaly in yeast extract supplemented RCV medium (4g Malic acid, 1g NH_4_Cl, 120 mg MgCl_2_, 75 mg CaCl_2_, 20 mg Sodium EDTA salt, 1mg Nicotinic acid, 10 mM KPO_4_ buffer, 1000 ml dH_2_O supplemented with 0.3g Yeast extract powder, *P* H 6.8 ±0.2). The culture was incubated at room temperature under continuous illumination for 6–8 days[29] in screw cap bottles. The full grown bacterial culture and the cell free extract were used for monitoring fluorescence properties.

### Spectroscopic Studies

Spectroscopic measurements of samples were obtained by scan method setting, absorption range from 250 nm to 1000 nm, at 1 nm bandwidth and 600 nm/min scan speed using Thermo Scientific *™* Evolution 300 UV- Vis spectrophotometer. The fluorescence properties of samples were studied with Photon Technology International (PTI) fluori-metric setup (Quantamaster *™*40). A 72 W Xenon lamp was used as an illumination source and the detection was preceded by passing the emitted beam through an optical chopper and emission monochromator. The slit width was set at 4 nm for both excitation and emission. For temperature dependent fluorescence measurement a software controlled Peltier was employed. The magnetic field application is illustrated in Fig. S1 of supplementary. A small square shaped magnet (field strength 295mT) was placed under the quartz cuvette, in the cuvette-holder of above mentioned fluorimetric set up to study the effect of static magnetic field (SMF). The temperature variation experiment in presence and absence of SMF was performed using this setup.

### Determination of kinetic parameters: Spectroscopic Data Analysis

Graphs were plotted considering the initial fluorescence amplification rate along Y axis and other variables such as temperature or concentration along the X axis. Change in fluorescence intensity (count) with per unit time was expressed by a first order kinetic parameter k (*s^−^*^1^). In case of amplification, k value is positive. A negative K value could be obtained if only bleaching occurs. To obtain k, only the initial rate was evaluated using first 20 seconds of data. Solution to the first order equation 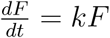, representing the rate of fluorescence intensity (F) change. Matlab (Mathworks USA) was used to obtain the best value of k. Multiple k values were determined from 5 independent experiments using log linear regression and this was followed by a box plot representation of data.

### Fluorescence Life Time Measurement

Fluorescence lifetime measurements were performed using 370nm sub-nanosecond pulsed LED source from HORIBA Scientific Single Photon Counting Controller: FluoroHub. Fluorescence Lifetime histogram was obtained using MicroTime 200, PicoQuant GmbH. Detail description of the instrument is given in the experimental set up section of Single Molecule Spectroscopy.

### Single Molecule Spectroscopy: Probe Molecule in Bulk Media and inside Cell

In the confocal microscope (MicroTime 200, PicoQuant GmbH), a picosecond diode laser with a base frequency of 40 MHz was used as excitation source. A water immersion objective (60X, 1.2 NA) was used to focus the excitation light (405 nm) from a pulsed diode laser (PDL 828-S “SEPIA II”, PicoQuant) onto sample kept on a coverslip. A dichroic mirror (Z405RDC,Chroma) and a filter (HQ430lp) were used to block the excitation light (405nm) along the path of fluorescence. The fluorescence was then focused through a pinhole (75 *μm*) onto the MPD detectors (Micro Photon Device). The emission spectra under a confocal microscope were recorded using an electron multiplying charge coupled device (EMCCD, ANDOR Technology) attached to a spectrograph (ANDOR Technology, Shamrock series). The spectrograph is attached to one of the ports of the PicoQuant Microtime 200 apparatus. In FCS, the autocorrelation function *G*(*τ*) of fluorescence intensity is defined as:

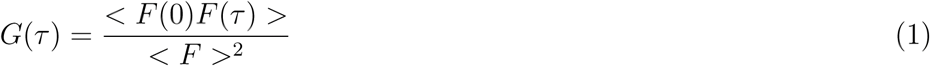

where *F* (0) and *F* (*τ*) denote the intensity of fluorescence at time 0 and at a lag time *τ*, respectively. The autocorrelation traces are fitted to a 3D-diffusion model[30, 31] having triplet contribution as,

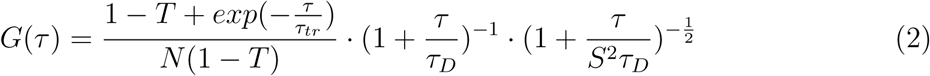

In the above equation, τ_*D*_ denotes the diffusion time in the confocal volume, τ_*tr*_ is the triplet lifetime of a dye molecule, T is the fraction of number of molecule in the triplet state, *τ* is the delay time and N represents the average number of molecules in the confocal volume. S (= *ω_z_/ω_xy_*) is the structure parameter of the excitation volume, *ω*_*z*_ and *ω_xy_* being the longitudinal and transverse radii respectively. S is calibrated using Rhodamine 6G (R6G) in water and is found to be 5. The diffusion constant *D_t_* of dye molecule was determined using the following equation,

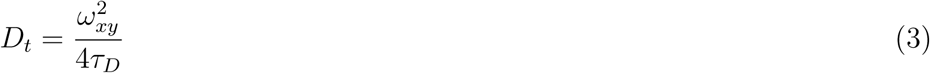

where, *ω_xy_* is the transverse radius (~ 320 nm) of the confocal volume (~ 0.9*f L*). The value of *ω_xy_* for our microscope was determined using R6G in water whose *D_t_* value (426 *μm*^2^*s^−^*^1^) is known[32]. From *D_t_*, we estimated the hydrodynamic radius *r_H_* of the fluorescent molecule applying Stokes-Einstein equation given by,

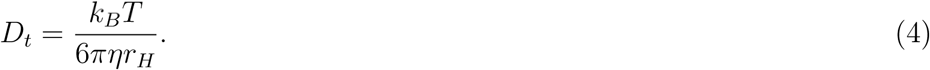

### Size Measurement of cell free extract by Dynamic Light Scattering (DLS)

400 *μ*L of sample was used for DLS measurement (Zetasizer Nano Series; Malvern Instruments, Malvern, UK) at 278K, 288K, 298K, 303K and 313K,respectively. Effect of a weak magnetic field on sample at 278K was studied by placing the magnet just below the cuvette during the measurement.

### Live Cell Imaging

Microscopic fluorescence study was carried out using bacterial biofilm [33] in an inverted confocal microscope (Olympus, FV 1000). A 405 nm laser was used as excitation source and fluorescence emission was observed between 570–670 nm. Time lapse imaging of the samples were acquired using continuous laser at 1 minute time interval for a period of 10 minutes. Additionally time trace emission from single bacterium and cell cluster were measured using the confocal microscope (MicroTime 200, PicoQuant GmbH).

## Results

### Spectroscopic analysis

Absorbance spectra of the cell free extract (see Fig. 1) show a soret band at 395 nm and Q band at 500 nm, 535 nm and 560 nm. The pattern of absorbance spectra indicates the presence of H and J band respectively near 400 nm and 535 nm (approximate ratio 7:1). The absorbance value of the sample shows a sustained increase with time of UV exposure.

**Figure 1.**
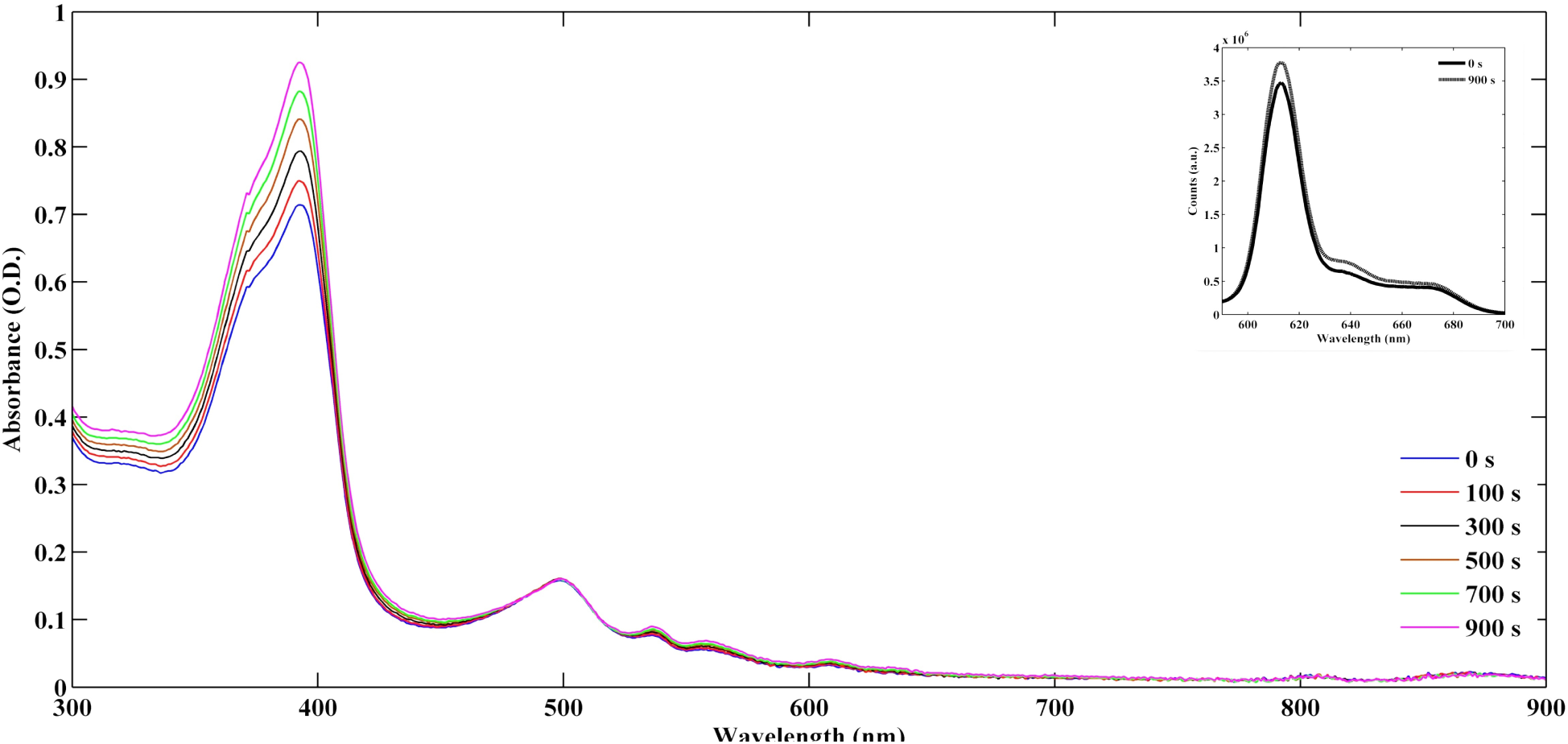
The absorbance spectra of the cell free extract were measured at particular time interval. Spectra shows gradual increase at 395 nm during prolonged excitation upto 900s. The inset shows initial (solid line) and final (dashed line) fluorescence spectra of the same sample indicating amplification in fluorescence.

The bacterial cells and the cell free extract showed fluorescence emission at 613 nm when excited at 395 nm. The time based fluorescence emission upon excitation at 395 nm is shown in supplementary figure S2. Time scan reveals a systematic increase in fluorescence emission intensity for both whole cells and the cell free extract. The rate of amplification in the two cases are however different. Organic extraction of the sample in acetone-methanol exhibited two peaks at 594 nm and 635 nm. Notably, the excitation is also red shifted to 411 nm and 403 nm for the respective emissions. The deconvolution of original emission peak through organic solvent perturbation (see Fig. S3 in supplementary information) into two peaks indicates the presence of two porphyrin derivatives that is coproporphyrin and Mg-protoporphyrin mono ethyl ester fluorescence [34]

### Molecular crowding and bistable emission

The cell free extract of bacterial culture shows transition in emitted fluorescence intensity over a threshold concentration. Fig. 2 reveals concentration dependent enhanced emission during a fluorescence time scan. Fig. 2 indicates that not only fluorescence intensity but also the amplification has a systematic dependence on the concentration; rate of emission (k) changing from positive to negative with dilution of the fluorophore concentration. One can find fluorescence emission intensity either rises or decreases with time depending on concentration of the fluorophore. Further insight of such concentration dependence is provided by a sigmoid signature in Fig. 2 where the initial (first 20 s) fluorescence emission rate has been studied.

**Figure 2.**
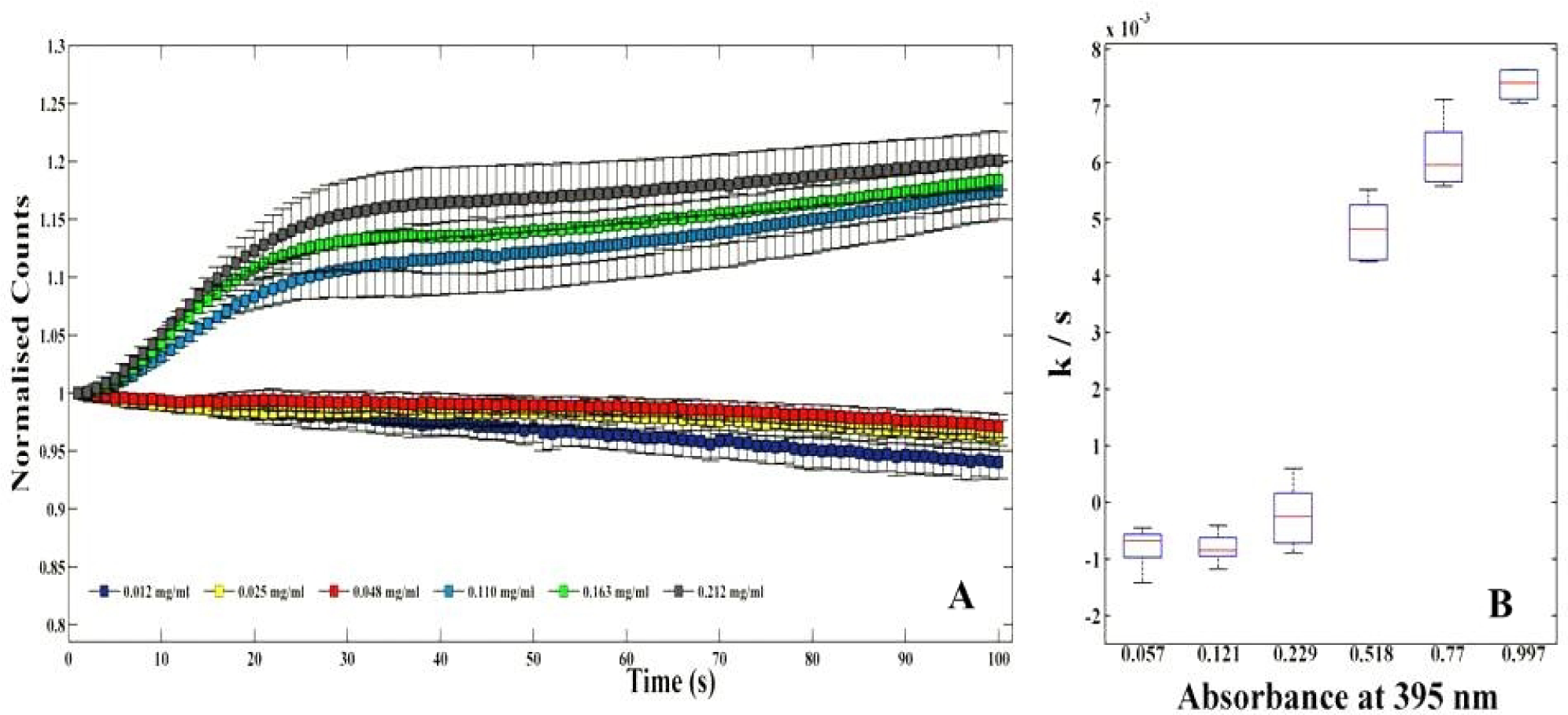
A. Time based fluorescence emission normalized with initial intensity along Y axis is plotted with concentration along X axis. Figure shows changing in fluorophore concentration switches the steady states from bleaching to enhancement of fluorescent emission. The bistability occurs between the critical concentration below 0.048 and above 0.11*mg·ml*^−1^; B. The initial rate (0–20s) of fluorescence emission(k) per unit time (s) is plotted with absorbance value at 395 nm assuming as a function of concentration of the samples. The graph shows sigmoid pattern in kinetics with increasing absorbance value.

### Temperature dependence and static magnetic field

The cell free extract of higher concentration (O.D. 0.1 at 395 nm) shows a systematic decrease in fluorescence counts at 278K in contrast to 298K (see upper panel of supplementary figure S4). But when the sample concentration is further diluted to ten times (O.D. 0.01 at 395 nm), shows no difference with temperature change. In lower panel of supplementary figure S4 the time based fluorescence of the above conditions are depicted. Fig. 3 illustrates the effect of Static Magnetic Field (SMF) on fluorescence property of the system. The amplification rate was found highest at 298K. But at low temperature (below 280K), the amplification process is attenuated (see upper panel of Fig. 3). The comparison of upper and lower panels of Fig. 3 clearly indicates that only at low temperature (278K), the amplification reappears even in presence of a low strength SMF.

**Figure 3.**
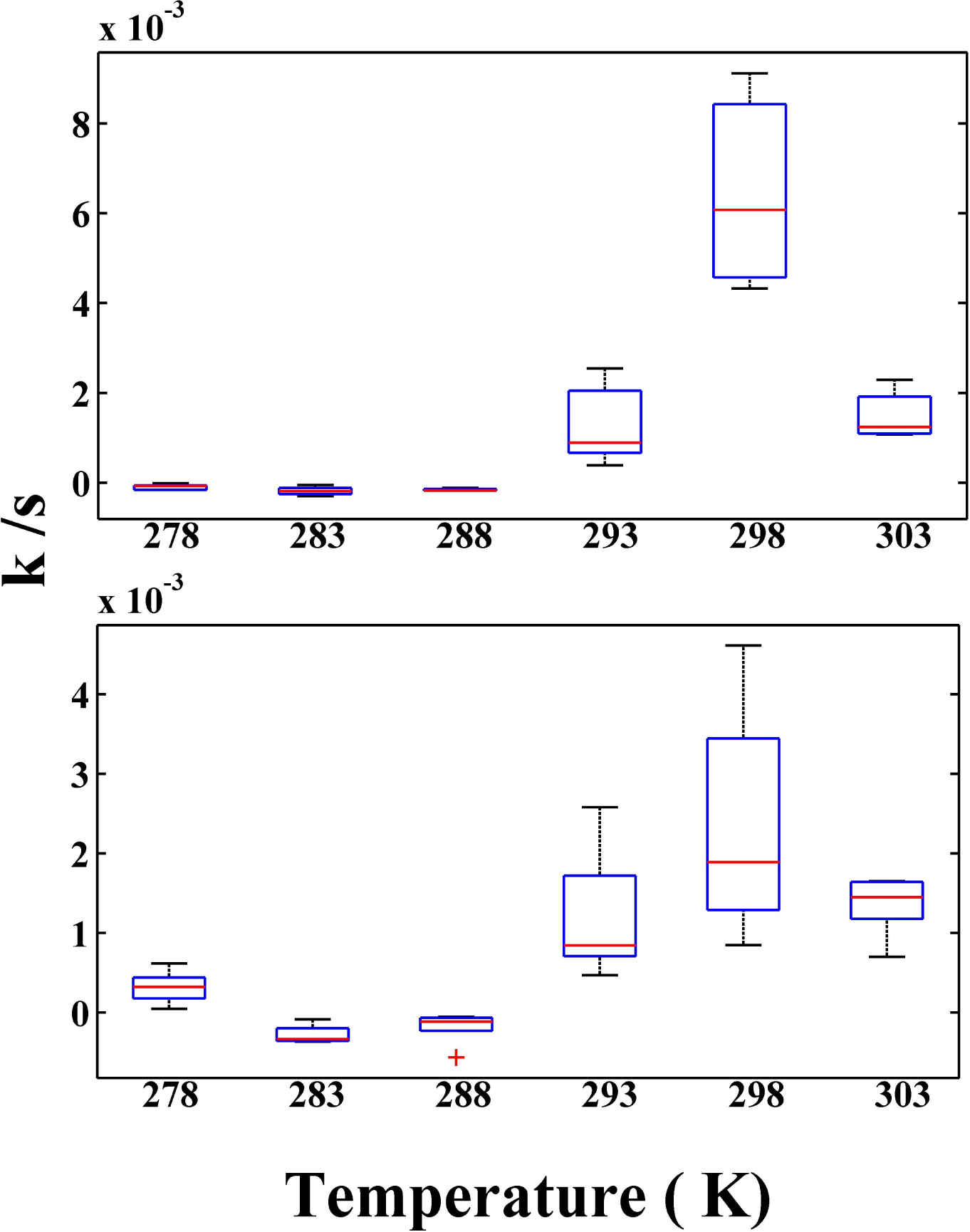
Fluorescence amplification rate at 613 nm (k s^−1^) is plotted with temperature in absence (upper panel) and presence (lower panel) of magnetic field. Figure shows in presence SMF the rate of amplification is positive at low temperature (278 K). Above 278 K the rate of enhancement is however lowered in presence of SMF.

### Fluorescence Life time

Life time studies revealed a few characteristics of the amplitude gain phenomenon. Dilution of sample shows that the life time value decreases with increasing order of dilution, higher dilutions have low life time values (see Table 1). The evaluation of life time values for the excreted fluorophore revealed order of 2 nanoseconds and 15 nanoseconds which indicate the formation of two species in excited states. So, the notable point is that during fluorescence emission only one species is found to show varying lifetime and another is not. In addition with life time data, the absorbance spectra obtained from flash photolysis experiment conforms presence of short living excited state singlets and long lived triplet states. The ration of the two has been found to alter upon time based excitation exposure.

**Table 1.**
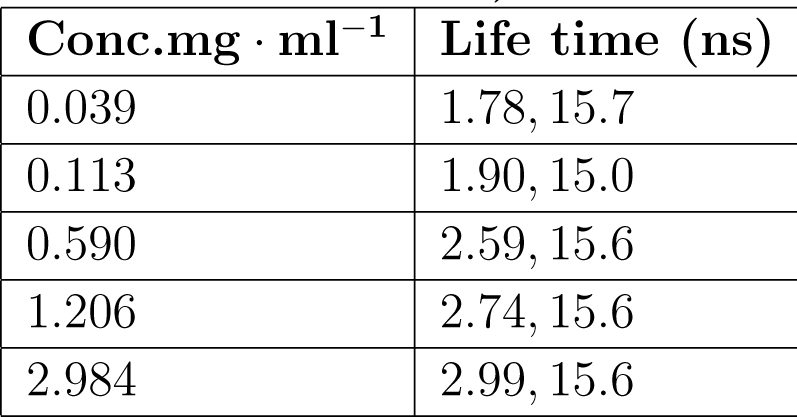
Life time values of the fluorophore at different concentrations (function of absorbance at 395 nm)

Life time variation with respect to fluorophore concentration.

### Fluorescence Correlation Spectroscopy

Fluorescence Correlation Spectroscopy with cell free fluorophores under such a condition where there is minor bleaching and also a small gain phase (see Fig. 4). In the Fig. 4 upper panel shows time trace emission spectra of the bacterial fluorophore and the lower panels show FCS decay patterns in two successive periods of time, one in which the bleaching dominates and the other in which there is a small gain over the steady state. Notably the FCS pattern and particularly the G(0) value remains identical in different time courses. The parameter values described in the table. 2 which additionally compares the FCS parameters for the sample and a control dye clearly indicates that the FCS parameters (explained in equations 1–4) practically remain unchanged in the two temporal phases.

**Figure 4.**
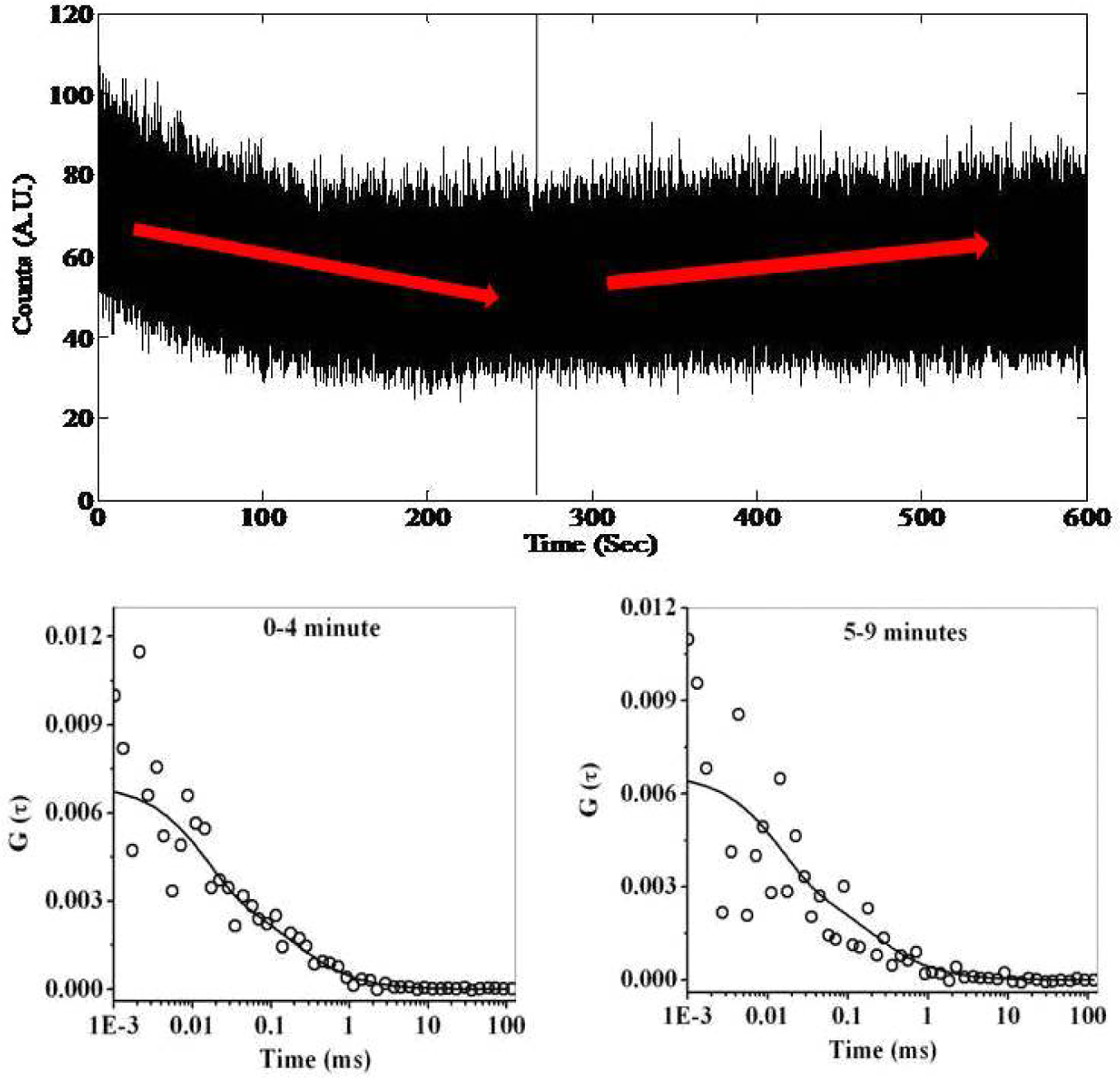
Upper panel shows time trace emission spectra of bacterial fluorophore of moderate concentration (0.1 mg/ml) where amplification occurs followed by initial bleaching. The lower panel shows FCS decay pattern of the bacterial fluorophore at two different time course. Initial phase showing dominance of bleaching and the later one for enhancement.

**Table 2.** The table shows different FCS (fluorescent correlation spectroscopy) parameter values (T,N,*τ_tr_,τ_D_,D_t_*) as defined in equation (2). The table compares parameter values for excreted bacterial fluorophores at two different time phased of FCS decay with the control dye coumarin 480(C480)

| System | | T | N | $\tau_{tr}$<br>(ms) | $\tau_D$<br>ms) | $D_t$<br>($\mu m^2 s^{-1}$) |
| --- | --- | --- | --- | --- | --- | --- |
| Sample(PC) | 0-4 Min | 0.5 | 300 | 0.015 | 0.17 | 150 |
|  | 5-9 Min | 0.38 | 315 | 0.015 | 0.18 | 142 |
| Control (C480) |  | 0.05 | 210 | 0.01 | 0.045 | 570 |

Comparison with the control dye (coumarin 480) is also depicted in the table. While the intensity pattern shows that there is an initial photo-bleach followed by recovery. This indicates that excitation of the fluorophore undergoes a photochemical transformation to form higher fluorescence quantum yield (brightness). This partially compensates for the bleach. The G(0) value and hence the number of molecules in the focal volume (N) of the FCS traces (Fig. 4) recorded at two different times (0–4 min and 5–9 min) are thus found to be identical in spite of bleaching or amplification.

However further insight with samples having varying fluorophore concentration shows a differential lifetime values of the excited triplets. The triplet life time value (standard deviation 10 %) is decreased with increasing degree of dilution. Whereas the cell free extract and the membrane fraction isolated from bacterial cells show more or less similar number of molecules in excited triplet state (see table. 3).

**Table 3.**
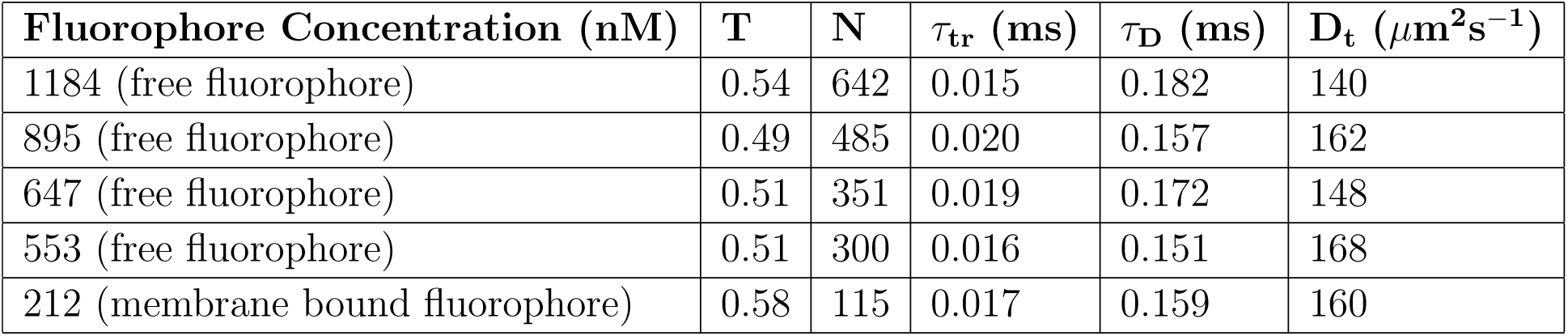
Characterization of cell free extract and membrane fraction through FCS

### Charecterization using Dynamic Light Scattering (DLS)

At low temperature (278K) the cell free extract shows abundance of small aggregates which is shifted to lager population size when exposed to a static magnetic field (see Fig. 5). DLS measurement of the sample shows temperature induced aggregation. In Fig. 6 when mean intensity was plotted against the hydrodynamic size, it was found that with increasing temperature there is a tendency of larger population formation with heterogeneous aggregate size. From Table 4 it is clearly observed that the Z average value of the sample is increasing with temperature. At 278K, the sample shows a increased Z average value in presence of SMF though the poly dispersity index is also increased. At 298K, 303K and 313K a very small population arises with respectively small hydrodynamic size which is absent in 278K, 288K and 313K (see Fig. 6 and Fig. 5).

**Figure 5.**
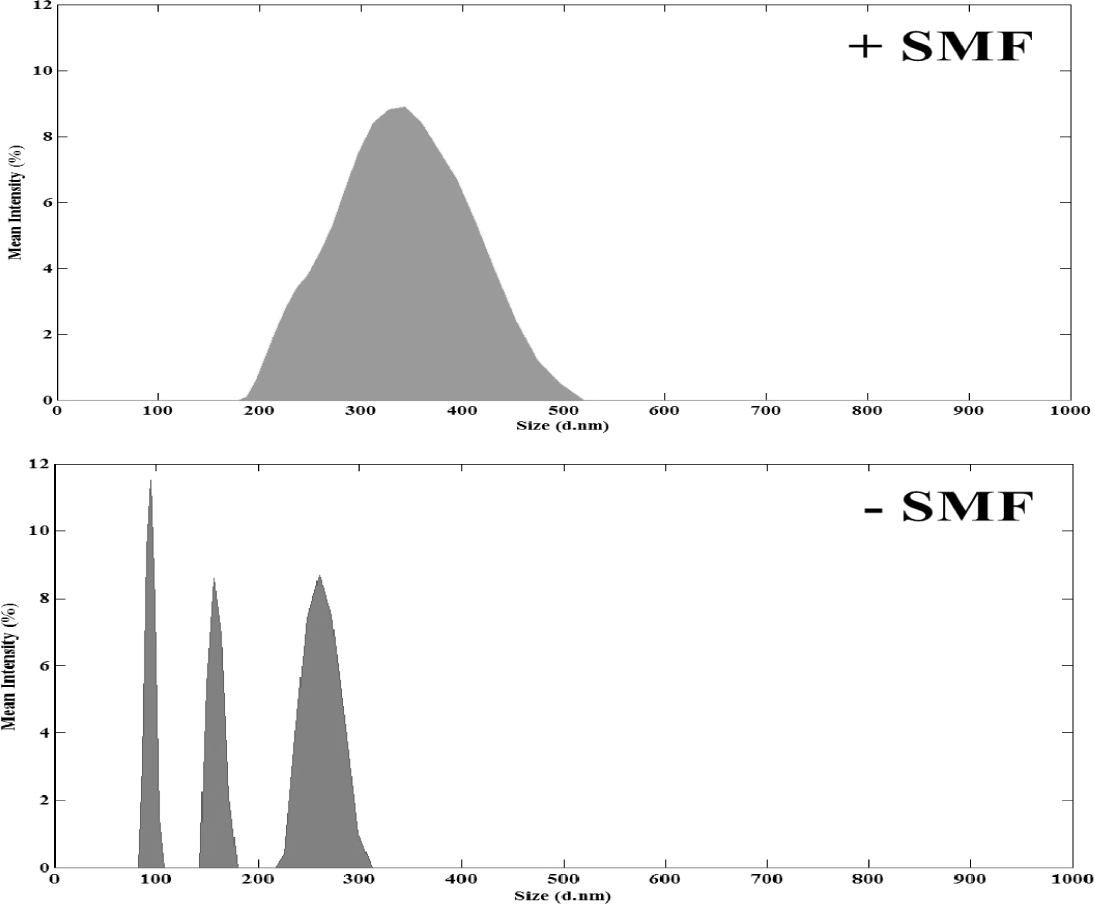
Dynamical light scattering measurement showing aggregate morphology in presence and absence of static magnetic field respectively at 278K. The sample tends to form larger aggregates in presence of SMF.

**Figure 6.**
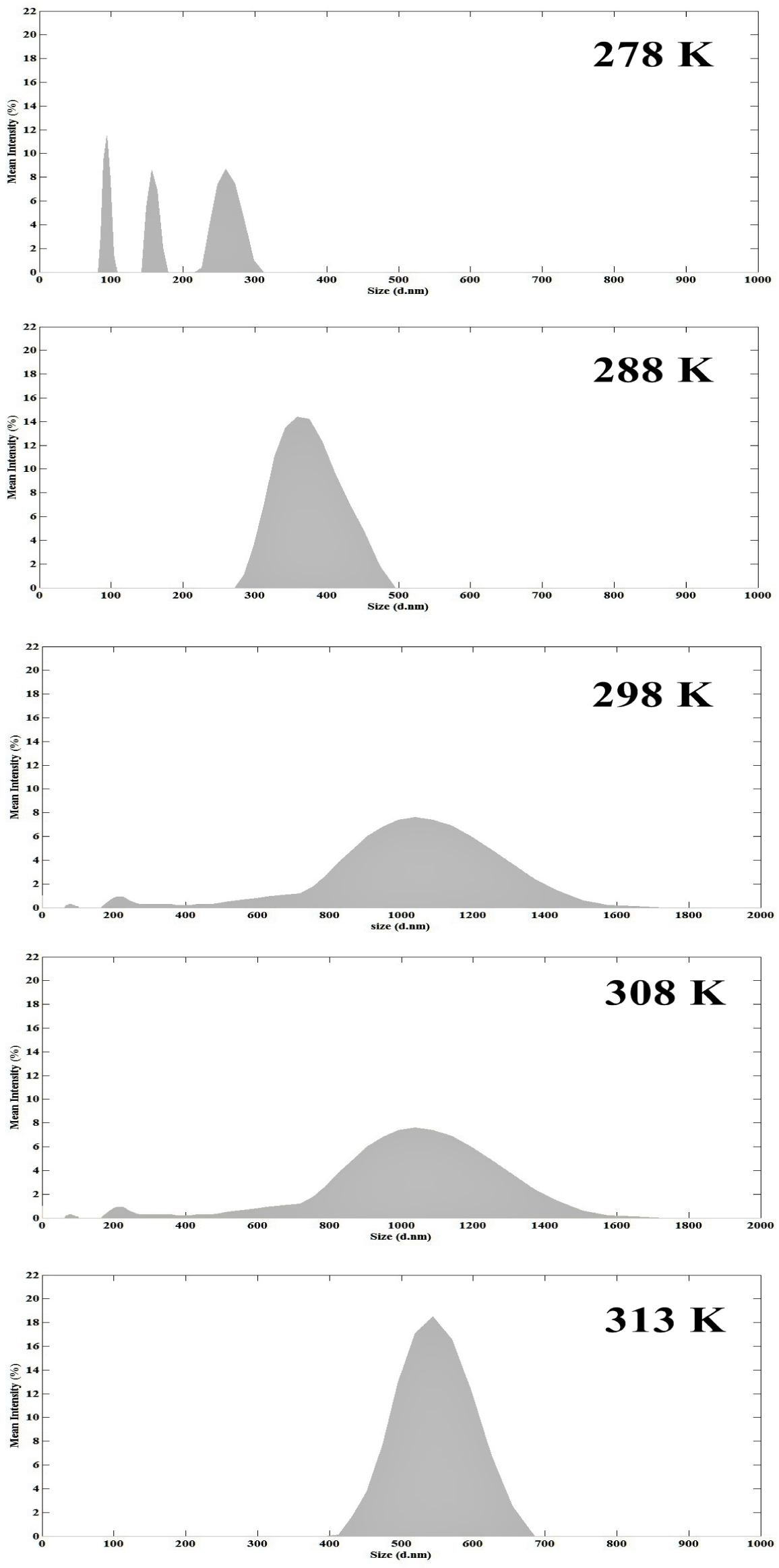
Temperature induced diversification of aggregate formation is presented in the figure. Higher temperature showing aggregates with higher hydrodynamic radius.

**Table 4.** T dependence of Polydispersity Index (PDI)

| Temperature | SMF (0.0285 T) | Z Average (d.nm) | Average PDI |
| --- | --- | --- | --- |
| 278K | — | $356.8 \pm 46.47$ | $\leq 1$ |
| 278K | + | $828 \pm 76.38$ | $< 0.5$ |
| 288K | — | $849.7 \pm 41.43$ | $\leq 1$ |
| 298K | — | $758.1 \pm 36.38$ | $\leq 0.5$ |
| 303K | — | $755.1 \pm 161.5$ | $< 1$ |
| 313K | — | $997.1 \pm 36.20$ | $\leq 1$ |

### Behavior of live cells

The microscopic data described in Fig. 7 reveals that whole bacterial cells showed enhanced fluorescence emission over a longer time period up to 10 minutes. The panels numbered 1,2,..10, corresponds to minutes of excitation exposure of the sample. When a FRAP (Fluorescence Recovery After Photo-bleaching) experiment was carried out with bacterial cells, the fluorescence emission intensity gradually increased instead of photo-bleaching. For further study the intensity of the incident laser (405 nm) beam was increased from 25%to 100% and the sample was exposed for one minute. The result is shown in the lower panel of Fig. 7. The indices A and B refer to per-excitation and post-excitation states of the sample. This time there was a dual effect, the central part showing a bleaching (see panel B of Fig. 7 and the peripheral region showing an intense fluorescence.

**Figure 7.**
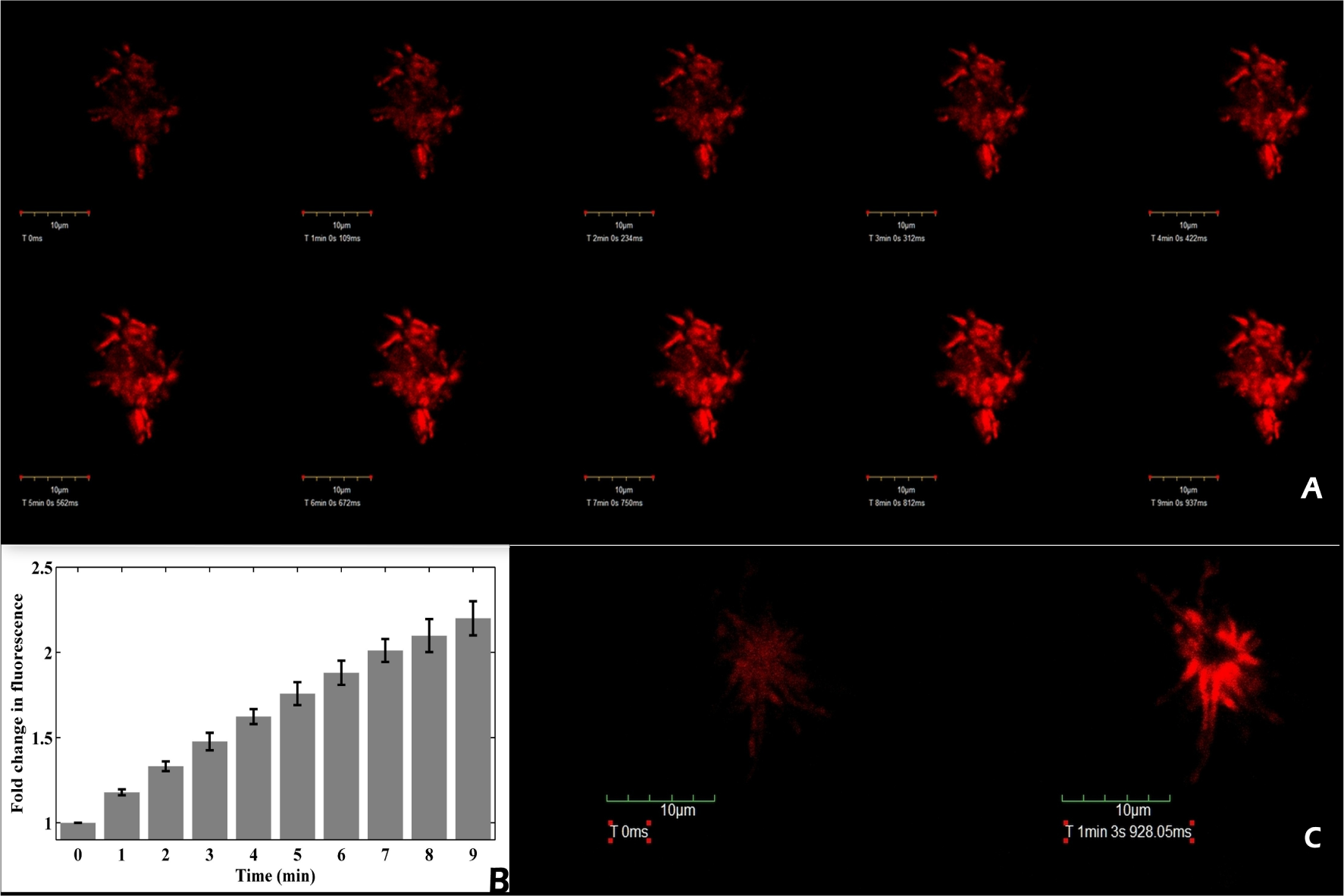
A. Gradual increase in fluorescence emission of bacterial cell cluster is observed. Cells were exposed to 405 nm laser (25%intensity) up to 9’, at 1’ interval starting from 0’; B. The relative increase in image intensity (with respect to 0 min, the error being evaluated using four indepndent ROIs); C. Image of a cell cluster exposed for 60 s to 405 nm laser (100%intensity), the initial image (left) after 60 seconds of laser exposure assumes the image form shown at the right (of the panel C). Whereas, the central zone of the right image is bleached due to intense laser exposure a more illuminated zone appears at some distance.

The temporal profile of intensity for single cell and cellular clusters are shown in Fig. 8. The gain in fluorescence with time in cellular clusters and the gradual decay of such intensity in single cells in the high resolution setup also conforms the preliminary observation reported in Fig. 7. The data is also a scaled up version of what we obtained in Fig. 2 which implies that as we increase the concentration, the amplification gain mode dominates.

**Figure 8.**
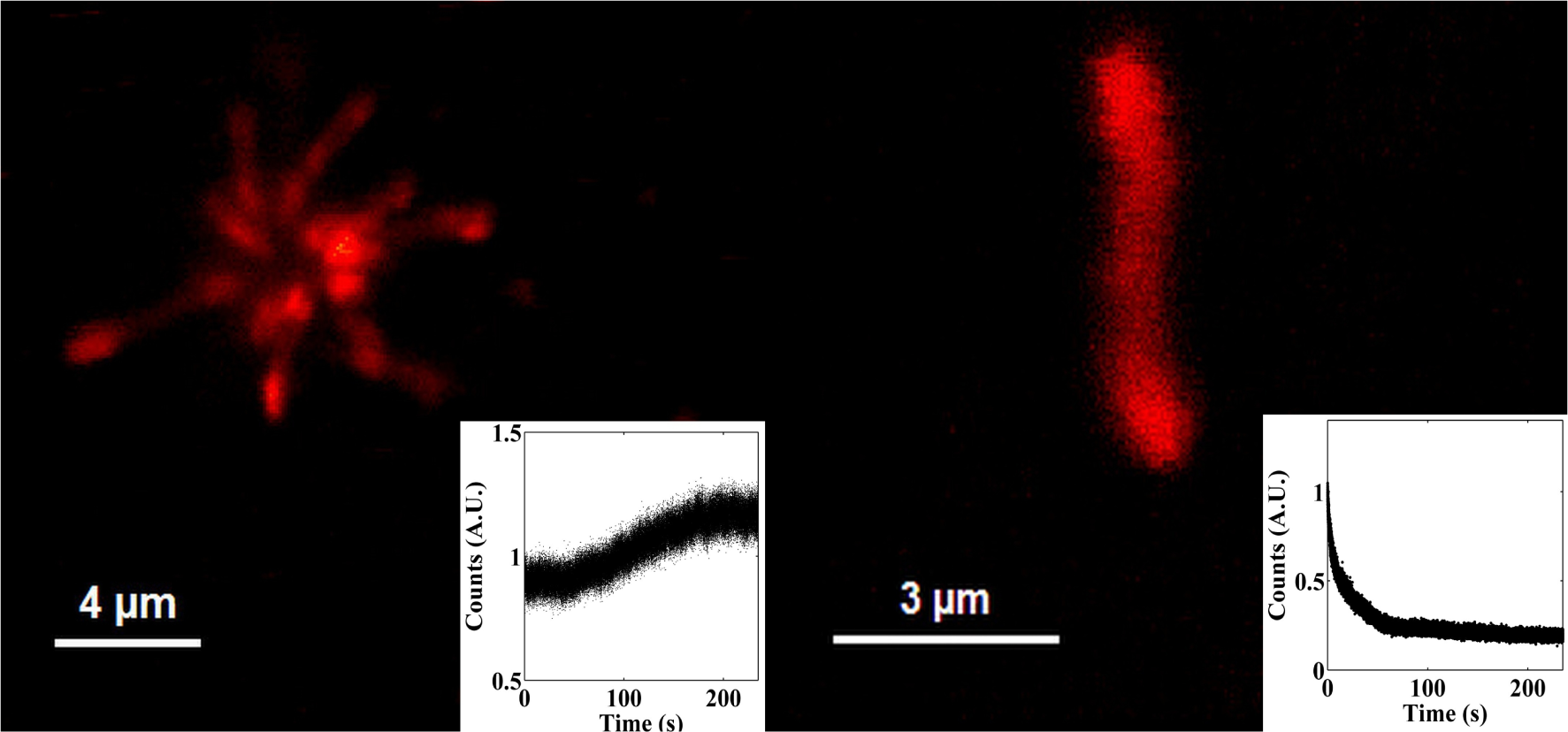
The difference in time trace emission of bacterial cell cluster and single cells has been depicted. The insets show increase in emission intensity for bacterial cell cluster and just opposite for single bacterium.

## Discussion

The paper reports long term incremental amplification of fluorescence intensity upon excitation in the UV region. The kinetics of increase in fluorescence is a switchable function of concentration, temperature and magnetic field. We may consider the following key aspects of the amplifying emission process.

- Amplification of fluorescence occurs at higher temperature and concentration
- Fluorescence quenches at low temperature and concentration
- Onset of amplification at low temperature (≤ 278*K*) in presence of SMF.
- The amplifying process studied with cell free extracts is also mimicked at cellular cell level, the cellular clusters showing the fluorescence amplifying behavior reverse being the scenario with single cells.

It appears that at a given state the overall change in fluorescence is result of a dynamic balance between the enhancement and bleaching processes, sign of k being determined by the dominating component. To explain the mechanism behind the amplified fluorescence emission we can have two following perspective views.

### Porphyrin Aggregation induced enhancement of Fluoresence

Photo excitation of porphyrin derivative leads to radical pair formation. However, the radical pair is known to be favored at lower temperature [26]. Incidentally in our case we get magnetic field effect only at lower temperature. Magnetic field affects only the population with finite spin as the singlet states are Zeeman insensitive (and would theefore remain insensitive to field). This is also reflected in the fluorescence emission kinetics. On the other hand, increase in local electrostatic potential of the surrounding medium upon prolong photo-exposure may facilitate the enrichment of singlets from fused triplets [35]. Thus the amplifying emission may be looked upon as gradual increase in the excited state population size.

When *Rhodobacter capsulatus* cells are grown under anaerobic or micro-aerophilic condition in presence of continuous light they do accumulate tetrapyrroles (Protoporphyrin, Mg-protoporphyrin derivatives) [34]. Porphyrins containing tetrapyrroles easily interact non-covalently because of the presence of pi bonds, commonly known as pi stacking and lead to self assembly [36, 37]. In higher concentration the fact is that the crowding of the photosynthetic molecule would favor photon induced self assembly formation which explain the concentration dependent rise in fluorescence. We may assume that a kinetically controlled self assembly gradually causes the enhancement process by accelerating the ground state energy transfer to the excited state. Whereas in low sample concentration, the number of monomeric porphyrins are diluted out hindering aggregation formation with time. While self assembly is known to be favored at low temperature [38], porphyrins undergo aggregate formation with increasing temperature [39]. Porphyrins with negatively charged peripheral groups easily aggregates in aqueous solution [40, 41]. In room temperature, porphyrins could coexist in monomeric and aggregated forms. Porphyrins when undergo aggregation, form J and H type of aggregates [42, 43] and they could remain in a mixture of the two [44] in room temperature. In Fig. 6 it could be clearly observed that there is highest population of various size of aggregates. This denotes that there is a dynamic equilibria between monomers and aggregates that should be the sole reason for fluorescence enhancement at room temperature in high concentration.

Magnetic field induced changes in the amplification pattern implies the role of stacked pi ring assemblies which are known to respond to static magnetic field[45]. Porphyrin J type aggregates are known to respond to magnetic field [46], when a magnetic field is applied they change their orientation and are aligned in more ordered structure [46]. Whereas the monomeric porphyrins are only sensitive to a magnetic field in low temperature due to lower thermal energy and much lower diffusion rate. Hence the aggregation formation should be favored in such condition. However in higher energy level a low strength magnetic field has very low effect due to increased thermal energy that supports our finding as we did not find any significant changes in time based fluorescence emission pattern in room temperature. However question may arise that why the overall emission rate is decreased in presence of SMF at room temperature. At normal temperature range due to very little effect of SMF the aggregation pattern will be realigned very slowly that should hamper the rate of enhanced fluorescence amplification and the process will be delayed.

### Thermodynamic Perspective : A Catathrope and its possible avoidance

The thermodynamic implications may be expressed by a simple expression for released thermal energy. We may consider a simple three state description of fluoresence (1 → 2 → 3) ∆*Q* for Stokes emission. Such emissions as observed in case of coral heating by GFP [47]. The opposite scenario namely cooling by anti-Stokes emission cooling[48] is also known.

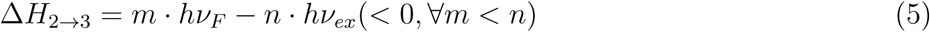

where m and n are number of photons involved in the excitation (1 → 2) and emission (3 → 1) process, the thermal energy being liberated during the vibrational relaxation of the higher excited state 2 to the lower excited state 3, the ground state being 1. The begative sign of the enthalpy change (heat produced) implies heating. Now if theer is amplification of fluoresence this will imply:

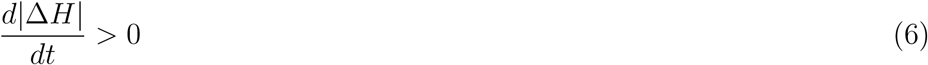

In other words, there will be uncontrolled heat generation. This can however be prevented if there is a compensatory heat transfer process via the porphyrin aggregate formation (that is favoured at higher temperature). The thermal catasthrope can be avoided if there is a vectorial transport of heat which in turn can have important ecological implications.

### Optical Communication by aggrgate formation

The confocal microscopic data in Fig. 7 implies that there is a gradual darkening of the irradiated zone and the higher illumination of the neighboring zone. A dynamic illuminated zone can be considered as an optical channel in microbial community. The amplifying emission may transfer the energy to the neighboring microbial layer (see Fig. 1). The figure shows that there is an absorbance ¿600nm, the regime that also has an emission when irradiated by UV. The serial energy transfer from lower to higher wavelength (in course of amplifiation) may be exploited by the cell

In parallel a chemically mediated the optical communication among microbes could also beactivated by photon induced aggregation of the porphyrin derivatives [49] and sensing of the same by neighbouring microbes. Porphyrin aggregation may thus constitute a component of photon sensing machinery for this class of photosynthetic bacteria.

### Bistability and Switching Behavior

Bistable fluorescence emission(Fig. 9) is another aspect of the work that deserves attention. The bistability in dynamical system is not a concurrent existence of two steady states. It rather implies possibility of transition from one steady state to another in response to variations of a critical parameter. One striking feature of Fig. 9 is that there are two routes of going to fluorescence -quenching state from the fluorescence-amplifying state, namely, thermal and magnetic. The co-operative change of the kinetic constant with respect to variations in concentration (Fig. 2) & temperature (Fig. 3) confirms onset of criticality in concentration or temperature dependent transitions.

**Figure 9.**
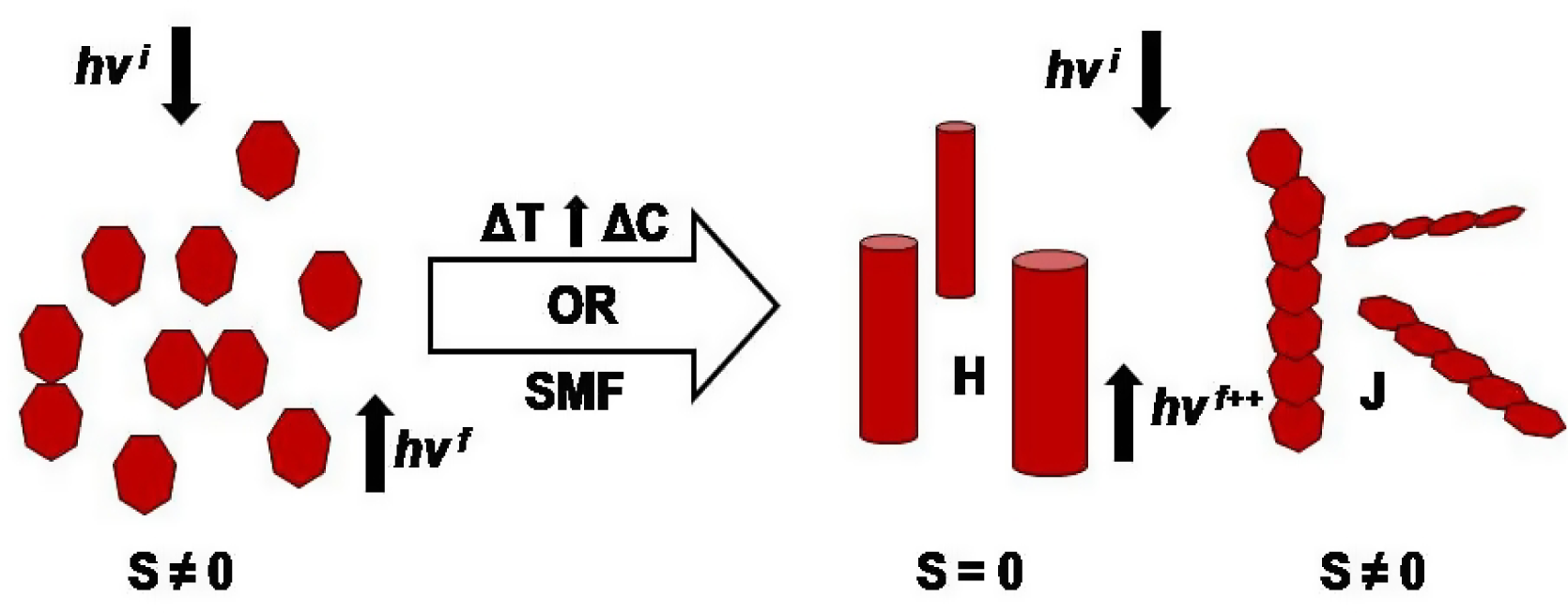
Schematic representation of aggregation induced fluorescence enhancement. T and C denotes temperature and concentration respectively. S represents spin states of excited states.

The striking feature of the reported bistability in *Rhodobacter capsulatus* is that more than one critical parameters lead to bifurcation of state from fluorescence quenching to amplification or *vice versa*.

### Low temperature magnetic sensitivity and the integrated narrative

Recently theoretical calculations have been made on porphyrin based nanowires [50]. The authors considered porphyrins as good candidates for molecular spintronics. Some of the porphyrin assemblies showed magnetic and some were nonmagnetic. What has been known so far is that sensitivity may partly be dependent on the supramolecular chirality of such aggregates [51]. Experimentally also it has been shown that porphyrin J aggregates with side by side assembly, may have uncompensated spin state. There is a notable difference in the aggregation pattern at low temperature as seen from Fig. 6 as compared to that prevailing at high temperature. Stacked monomers forming H aggregates with neutralized spin state may show such property. This is possible if the magnetic moments of the monomers neutrilize to make overall zero spin state. The additional perspective which we think will be a promising future work is to check whether the magnetic insensitive assembly has higher qunatum yield as compared to the magnetically sensitive assembly, the later apperaing at lower temperature.

## Conclusion

The bistable behavior is shown for a photosynthetic population. Most of the previous reports on bistability were based on expression of single gene or proteins. It is also intriguing to note that the principle of population inversion on which the lasers are made is exploited by one of the most ancient of life forms. The concentration, temperature and magnetic field sensitive switches seem to suggest that logical system can prevail in the population. Photons one one hand are translated to metabolic route on the other hand is exploited at the intercellular level for cell cell communication in a logical way. No amplification of fluoresence when there is an incident UV irradiation may be a code by which the population senses low temperature. The magnetic field induced reversal of amplification can again serve as a vectorial code to the microbe community.

## Acknowledgment

We thank Department of Biotechnology, Govt. of India for supporting the research (grant number BT/PR3957/NNT/28/659/2013) and Department of science and Technology IRPHA project. We thank Dr. Shibsekhar Roy, Dr.Swagata Banerjee and Dr. Jaydeep Bhattacharya for helpful suggestions and Ms. Boni Haldar, Ms. Puja Biswas (DBT IPLS) for technical assistance in confocal and AFM experiments. We also thank Prof. M.Bhattacharya for providing the life time measurement facility.

